# Anionic lipids regulate PLCβ membrane recruitment

**DOI:** 10.64898/2026.07.20.739664

**Authors:** Makayla A. Crawford, Marie B. Bennett, Yu Qin, Kristin E. Cano, Taylor G. Gonzalez, Rahul Mojidra, Maria E. Falzone

## Abstract

Phospholipase C-β (PLCβ) enzymes are essential effectors of G protein-coupled receptor signaling that hydrolyze phosphatidylinositol 4,5-bisphosphate (PIP2) at the plasma membrane to generate the second messengers inositol trisphosphate and diacylglycerol. PLCβs play essential roles in diverse physiological processes, including cardiac and neuronal function, macrophage activation, and the pathogenesis of diseases such as hypertrophic cardiomyopathy. PLCβ enzymes are unique in that they are aqueous-soluble and must partition onto the membrane surface to access their substrate, making membrane association a critical regulatory step. For example, we recently demonstrated that Gβγ activates PLCβ by membrane recruitment and orientation of the catalytic core on the membrane surface. Although PLCβ membrane recruitment is required for function, the molecular determinants governing this process remain incompletely understood. Using a quantitative membrane partitioning assay, we show that robust membrane association of PLCβ requires anionic phospholipids, whereas polar phospholipids cannot substitute. Membrane partitioning exhibits a steep dependence on anionic lipid abundance, which is mediated by electrostatic interactions between negatively charged lipids and basic residues within the distal C-terminal domain of PLCβ. We further demonstrate that anionic lipids cooperate with Gβγ to regulate PLCβ membrane recruitment, such that the magnitude of Gβγ-dependent recruitment is dictated by membrane anionic lipid content. These findings reconcile previous discrepancies regarding Gβγ-mediated PLCβ membrane recruitment and establish membrane electrostatics as a key regulatory input that integrates lipid composition with G protein signaling to regulate PLCβ activity.

**Summary:** Membrane association is a crucial component of PLCβ function and regulation, yet the mechanisms governing this process are poorly understood. The authors demonstrate that anionic lipids drive PLCβ membrane recruitment and influence Gβγ-dependent activation, revealing how membrane composition shapes cellular signal transduction.

## Introduction

PLCβ enzymes are a canonical downstream target of G protein-coupled receptor (GPCR) signaling via direct activation by the G proteins Gβγ and Gα_q_(Kadamur and Ross, 2013; Lyon and Tesmer, 2013; Ubeysinghe et al., 2023). Upon activation, PLCβ enzymes cleave phosphatidylinositol 4,5-bisphosphate (PIP2) from the plasma membrane, producing inositol triphosphate (IP3) and diacylglycerol (DAG), which elevate intracellular Ca^2+^ and activate protein kinase C, respectively (Rodnight, 1956; Kemp et al., 1961). PLCβ function is essential for a myriad of cellular processes including cardiac and neuronal function as well as macrophage activation. Accordingly, dysregulation of these enzymes leads to human diseases, including neurological disorders, cancer, and cardiac hypertrophy (Kadamur and Ross, 2013; Lyon and Tesmer, 2013; Ubeysinghe et al., 2023). Thus, it is critical to understand the molecular mechanisms underlying their function and regulation under normal physiological conditions and dysregulation in human disease. Substantial progress has been made in characterizing PLCβ function, however aspects of their regulation remain poorly understood due to their intricate relationship with the membrane. PLCβ enzymes are unique in that they are aqueous-soluble but must partition onto the membrane surface to access their lipid substrate, which allows these enzymes to be regulated at both the catalytic step and the partitioning step. We recently showed that the G proteins Gβγ and Gα_q_ activate PLCβ enzymes utilizing both of these mechanisms, Gβγ by membrane recruitment and orientation of the catalytic core on the membrane surface and Gα_q_ by increasing the catalytic rate constant (Falzone and MacKinnon, 2023b; Falzone and MacKinnon, 2023a; Falzone et al., 2026).

Given the importance of PLCβ membrane association for function and regulation, it is essential to understand the drivers of this process and their underlying molecular mechanisms. Membrane association of PLCβ enzymes involves direct interactions with phospholipids, suggesting that phospholipid chemical properties may influence this association. Further, the substrate lipid PIP2 is not required for membrane association, indicating additional PLCβ contacts with non-substrate phospholipids underlie this association (Romoser et al., 1996; Runnels et al., 1996; Jenco et al., 1997; Falzone and MacKinnon, 2023b; Falzone and MacKinnon, 2023a). There are numerous types of lipids in the plasma membrane, but the chemical properties of the head groups with which PLCβ enzymes interact can be grouped into non-polar, polar, or negatively charged (**Fig. 1A**) (Robinson et al., 2019; Levental and Lyman, 2023). At rest, the plasma membrane is asymmetric with charged and polar lipids concentrated on the inner leaflet. This asymmetry generates a net negative charge on the inner leaflet of the plasma membrane where PLCβ interacts (**Fig. 1A**). Additionally, phospholipid remodeling is a frequent outcome of signaling cascades, which would facilitate additional layers of PLCβ regulation to fine-tune their localization and function for the needs of the cell (Robinson et al., 2019; Levental and Lyman, 2023). Accordingly, we hypothesize that PLCβ membrane association is dependent on non-substrate phospholipid chemical properties.

**Figure 1:**
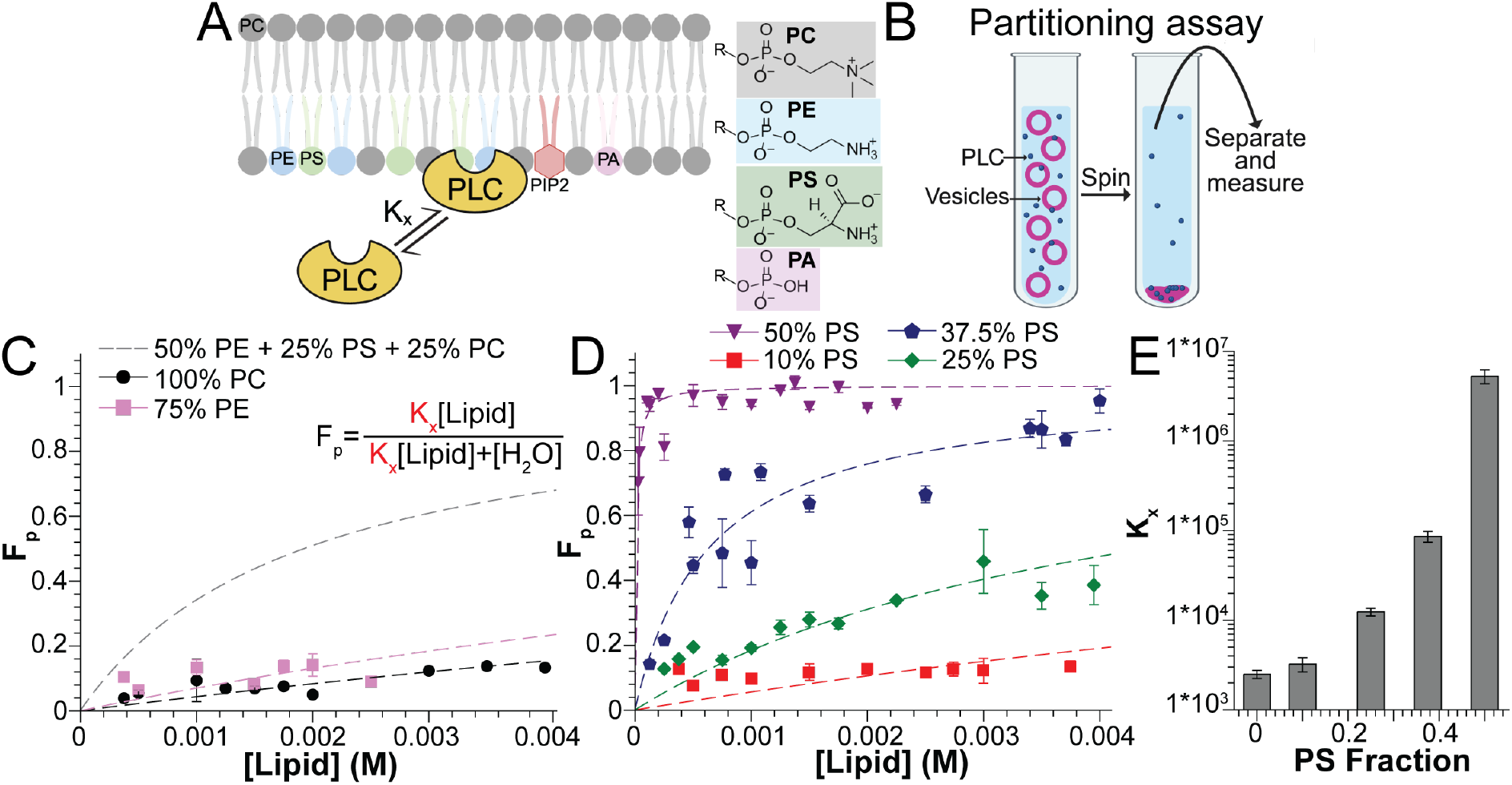
PLCβ membrane partitioning is dependent on PS content. **A**: Lipid distribution in the plasma membrane with lipid headgroup chemical structures for PC, PE, PS, and PA shown as examples. **B**: Cartoon for membrane partitioning assay. **C-D**: Membrane partitioning curves for PLCβ in 100% PC lipids (black) and 75% PE lipids (pink) (C) or increasing proportions of PS (D). Symbols are individual lipid concentrations across at least 2 PLCβ concentrations and dashed lines are fits to Equation 2 with values reported in Table 1. Error bars are range of mean. The fit for PLCβ partitioning in the presence of 2PE:1PC:1PS lipids (Falzone and MacKinnon, 2023a) is shown are a gray dashed line in C for comparison. **E**: Comparison of K_x_ as a function of PS content. Error bars are the error from the fit to determine K_x_.

Previous attempts to address this question have been contradictory, with some reports suggesting that phospholipid composition has no effect on PLCβ membrane association, while some suggest negatively charged lipids are essential for this association (Romoser et al., 1996; Runnels et al., 1996; Jenco et al., 1997). We have revisited this question with an updated framework for PLCβ function and regulation as well as technical approaches. Here, we demonstrate that robust membrane association of PLCβ enzymes requires negatively charged lipids, with specific electrostatic PLCβ-phospholipid interactions mediated by arginine and lysine residues in the distal C-terminal domain (dCTD) of the protein. Further, our results demonstrate that negatively charged lipids cooperate with Gβγ to fine-tune PLCβ membrane localization and function. These studies clarify discrepancies in previous studies regarding Gβγ-induced enhancement of membrane association (Romoser et al., 1996; Runnels et al., 1996; Jenco et al., 1997; Fisher et al., 2020; Fisher et al., 2026). Together, our findings establish membrane electrostatics as a fundamental layer of PLCβ regulation, providing a mechanism by which dynamic changes in lipid composition and G protein signaling are integrated to tune PLCβ activity in space and time.

**Table 1:** All measured K_x_ values with the error from the fit to Equation 2.

| Lipid Composition | Salt Concentration (mM) | K<sub>x</sub> (from fit) |
| --- | --- | --- |
| 50% PE + 25% PS + 25% PC | 150 | $2.9 \cdot 10^4 \pm 1.8 \cdot 10^3$ (7) |
| 100% PC | 150 | $2.5 \cdot 10^3 \pm 2.6 \cdot 10^2$ |
| 10% PS + 90% PC | 150 | $3.2 \cdot 10^3 \pm 5.8 \cdot 10^2$ |
| 25% PS + 75% PC | 150 | $1.2 \cdot 10^4 \pm 1.2 \cdot 10^3$ |
| 37.5% PS + 62.5% PC | 150 | $8.6 \cdot 10^4 \pm 1.2 \cdot 10^4$ |
| 50% PS + 50% PC | 150 | $5.2 \cdot 10^6 \pm 9.4 \cdot 10^5$ |
| 75% PE + 25% PC | 150 | $4.1 \cdot 10^3 \pm 6.7 \cdot 10^2$ |
| 25% PG + 75% PC | 150 | $5.0 \cdot 10^3 \pm 7.2 \cdot 10^2$ |
| 37.5% PG + 62.5% PC | 150 | $9.3 \cdot 10^5 \pm 1.2 \cdot 10^5$ |
| 50% PG + 50% PC | 150 | $1.0 \cdot 10^6 \pm 1.6 \cdot 10^5$ |
| 37.5% PS + 62.5% PC | 50 | N/A |
| 37.5% PS + 62.5% PC | 250 | $1.2 \cdot 10^4 \pm 9.5 \cdot 10^2$ |
| 37.5% PS + 62.5% PC | 500 | $3.6 \cdot 10^3 \pm 5.4 \cdot 10^2$ |
| 25% PS + 75% PC | 50 | $9.4 \cdot 10^5 \pm 1.3 \cdot 10^5$ |
| 25% PS + 75% PC | 250 | $6.2 \cdot 10^3 \pm 8.6 \cdot 10^2$ |
| 37.5% PS + 62.5% PC (ΔdCTD) | 150 | $1.0 \cdot 10^3 \pm 2.2 \cdot 10^2$ |
| 50% PS + 50% PC (ΔdCTD) | 150 | $1.0 \cdot 10^3 \pm 1.2 \cdot 10^2$ |
| 37.5% PS + 62.5% PC (24RK-A) | 150 | $2.5 \cdot 10^3 \pm 2.6 \cdot 10^2$ |
| 50% PS + 50% PC (24RK-A) | 150 | $1.7 \cdot 10^3 \pm 2.1 \cdot 10^2$ |
| 100% PC + G$\beta\gamma$ | 150 | $1.7 \cdot 10^4 \pm 1.8 \cdot 10^3$ |
| 10% PS + 90% PC + G$\beta\gamma$ | 150 | $1.1 \cdot 10^4 \pm 9.4 \cdot 10^2$ |
| 25% PS + 75% PC + G$\beta\gamma$ | 150 | $2.2 \cdot 10^5 \pm 1.7 \cdot 10^4$ |

## Results

### Robust PLCβ membrane partitioning requires anionic lipids

We previously utilized a rigorous assay for quantifying PLCβ membrane association using the framework of partitioning between two immiscible solvents, the aqueous solution and the hydrophobic membrane, rather than discrete ligand binding (White et al., 1998; Falzone and MacKinnon, 2023a). This framework accounts for the dynamic nature of membrane association of peripheral membrane proteins, which typically does not occur through fixed binding sites or stoichiometry (White et al., 1998). The equilibrium of PLCβ between the aqueous solution and the membrane is dictated by the partition coefficient K_x_, which is equal to the ratio of the mole fraction of PLCβ on the membrane to the mole fraction of PLCβ in the aqueous solution (Equation 1).

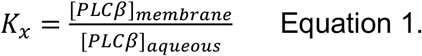

To measure K_x_, we employed a vesicle pelleting assay, enabling the measurement of the fraction of PLCβ partitioned (F_p_) across a wide range of lipid concentrations (**Fig. 1B**). Upon rearrangement, Equation 1 gives rise to Equation 2.

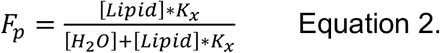

Accordingly, the F_p_ as a function of lipid concentration can be fit to determine K_x_ (White et al., 1998; Falzone and MacKinnon, 2023a). We previously determined the K_x_ for PLCβ3 to be 2.9*10^4^ in a mixed lipid composition comprised of 2PE:1PC:1PS (wt:wt:wt) or 50% polar, 25% non-polar, and 25% negatively charged lipids by mass (Falzone and MacKinnon, 2023a) (**Fig. 1C**). In this composition, partitioning is such that even at high lipid concentrations (~4 mM), only ~70% of the PLCβ protein is membrane-associated. This is consistent with recent cellular experiments demonstrating that most PLCβ3 is intracellular at rest (Senarath et al., 2025; Falzone et al., 2026; Fisher et al., 2026). To investigate the role of phospholipid headgroup chemistry in regulating PLCβ membrane partitioning, we first measured the K_x_ in the context of 100% non-polar lipids (100% PC). Under these conditions, K_x_ is reduced by ~10-fold to 2.5*10^3^, suggesting that anionic or polar lipids are required for robust membrane partitioning (**Fig. 1C, S1A, Table 1**). To evaluate this hypothesis, we measured K_x_ in the context of differing proportions of polar and anionic lipids dictated by mass. In the presence of 75% polar lipids (PE), K_x_ was not significantly different from that in 100% PC (4.1*10^3^), suggesting that polar lipids alone are not sufficient to mediate robust partitioning (**Fig. 1C, S1B, Table 1**). To assess the requirement of anionic lipids for partitioning, we measured K_x_ across increasing PS concentrations. 10% PS did not significantly increase K_x_ (3.2*10^3^), however, 25% PS increased K_x_ ~5-fold to 1.2*10^4^. 37.5% PS increased K_x_ an additional ~7-fold to 8.6*10^4^, and 50% PS increased K_x_ an additional ~60-fold to 5.3*10^6^ (**Fig. 1D-E, S1C-F, Table 1**). Further, K_x_ in the presence of 50% PS is ~2,000-fold higher than in 100% PC, illustrating that PS is a significant driver of PLCβ membrane partitioning in the physiological content range (**Fig. 1D-E, S1C-F, Table 1**) (Platre and Jaillais, 2017; Levental and Lyman, 2023). Interestingly, partitioning in the presence of 25% PS differs by ~2-fold in the presence and absence of PE lipids, suggesting that polar lipids can contribute to PLCβ membrane association in the presence of anionic lipids but are not sufficient independently (**Fig. 1C-D, Table 1**).

To connect the PLCβ membrane recruitment capacity of PS to PIP2 hydrolysis, we measured PLCβ lipase activity with differing proportions of PS using a fluorescence-based assay with the probe XY-69 (Huang et al., 2018; Carr et al., 2021), which allowed for full control of the lipid composition. The fluorescence of XY-69 is quenched in the intact molecule and upon hydrolysis, the quencher is released resulting in an increase in fluorescence, thus reporting on lipase activity (**Fig. 2A**). Hydrolysis is very low in the absence of PS but increases significantly with increasing PS content (**Fig. 2B-C, S1G-I**). The hydrolysis curves in the presence of 25% and 50% PS are well fit by a double exponential function, whereas the curves in the presence of 100% PC are better fit by a combined exponential and linear function (see Materials and Methods). Upon comparison of the first exponential component across all 3 conditions, the amplitudes increase significantly with increasing PS content, and the time constants (τ) decrease significantly, indicating higher activity (**Fig. 2C, S1G**). These observations are consistent with different amounts of PLCβ present on the membrane as demonstrated in our partitioning experiments (**Fig. 1D-E**), consistent with PS serving as a significant driver of PLCβ membrane recruitment.

**Figure 2:**
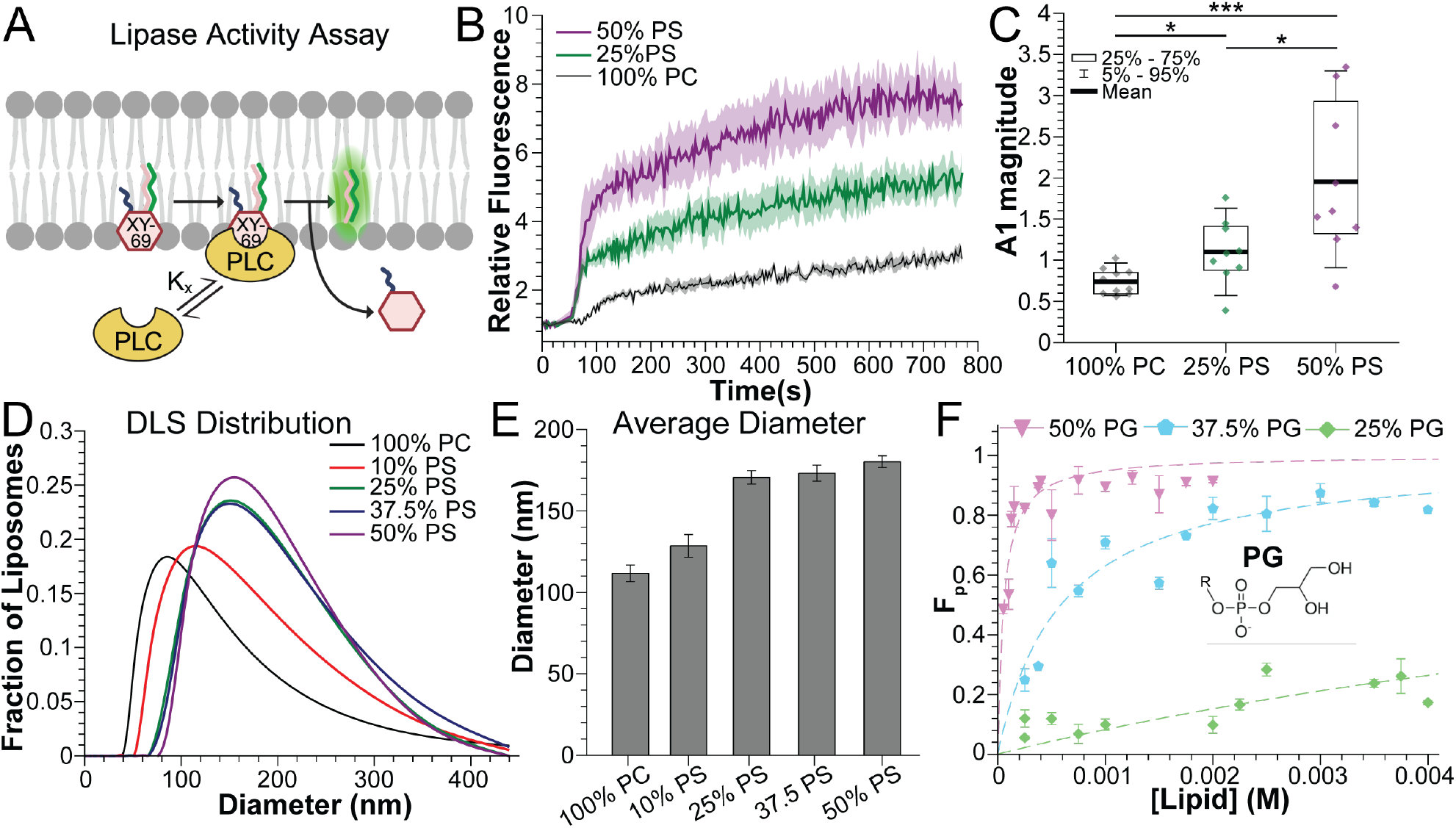
Anionic lipids are required for robust PLCβ membrane partitioning. **A**: Cartoon for XY-69 fluorescence assay (Huang et al., 2018; Carr et al., 2021). **B**: PLCβ activity on 500 μM of liposomes comprised of 100% PC (black), 25% PS (green) or 50% PS (purple) containing 0.1 mol% XY-69 and 2 mol% PIP2. Curves are an average of 3 wells and error is SEM. **C**: Comparison of the amplitude of the first exponential component of the fit for each condition (Amplitude 1). Diamonds represent each replicate. *: 0.05>p>0.005. **: 0.005>p>0.0005. ***: p<0.0005. **D**: Diameter distribution from representative DLS measurements across varying PS contents. **E**: Average vesicle diameter reported from DLS measurements. Error bars are SEM. **F**: Membrane partitioning curves for PLCβ in the presence of 25% (green), 37.5% (blue), or 50% (purple) PG. Symbols are individual lipid concentrations across at least 2 PLCβ concentrations and dashed lines are fits to Equation 2 with values reported in Table 1. Error bars are range of mean.

Further, to ensure that our observations are due to the change in the proportion of PS rather than changes in size or curvature of the vesicles, we used dynamic light scattering (DLS) to measure the vesicle size distribution as a function of PS content (**Fig. 2D-E**) (Khan et al., 2022). The size distributions shift to slightly larger sizes as anionic lipid content is increased up to 25%, with the largest average difference of ~70 nm between 100% PC and 50% PS. However, the distributions for 25%, 37.5% and 50% PS are virtually identical, despite the ~400-fold and ~60-fold difference in PLCβ partitioning in these conditions (**Fig. 1D-E**). Accordingly, it is unlikely that the <2-fold difference in average vesicle size underlies the ~2,000-fold increase in membrane association observed between 100% PC and 50% PS, highlighting a specific role of PS in PLCβ membrane recruitment.

Finally, we investigated if the observed PS-mediated membrane recruitment is specific to PS or if any anionic lipid can mediate partitioning using phosphatidylglycerol (PG), which also carries a charge of -1 (**Fig. 2F**). Increasing concentrations of PG lead to similar increases in K_x_, with 25% increasing K_x_ ~2-fold compared to PC alone to 5.0*10^3^, 37.5% PG increasing K_x_ another ~20-fold to 9.3*10^4^, and 50% PG increasing K_x_ an additional 10-fold to 1.0*10^6^ (**Fig. 2F, S2, Table 1**). Further, K_x_ is similar across the same proportions of PG and PS (**Fig. S2A**), indicating that the negative charge rather than a specific lipid species is important for robust membrane association of PLCβ.

### Electrostatic interactions between the PLCβ dCTD and anionic lipids underlie partitioning

Given the strong dependence of partitioning on anionic lipids, we hypothesized that electrostatic interactions between PLCβ and the membrane are responsible for robust PLCβ membrane association. To address this hypothesis, we modulated the strength of these electrostatic interactions by varying the salt concentration in our partitioning experiments. In agreement with our hypothesis, reducing the salt concentration from 150 mM KCl to 50 mM KCl resulted in a robust increase in membrane partitioning in both 25% and 37.5% PS (**Fig. 3A, S3, Table 1**). In the presence of 25% PS, K_x_ increased ~100-fold to 9.4*10^5^, however, in the presence of 37.5% PS, partitioning was increased such that we were unable to accurately fit the data to Equation 2, as PLCβ was fully partitioned at 50 µM lipid, which is the lowest concentration we can reliably measure. Further, increasing the salt concentration from 150 mM to 250 mM or 500 mM significantly reduced partitioning in the context of both 37.5% PS (~7-fold for 250 mM, 24-fold for 500 mM) and 25% PS (~2-fold for 250 mM) (**Fig. 3A, S3, Table 1**). These observations support the hypothesis that electrostatic interactions between PLCβ and anionic lipids underlie robust membrane association (**Fig. 3B**).

**Figure 3:**
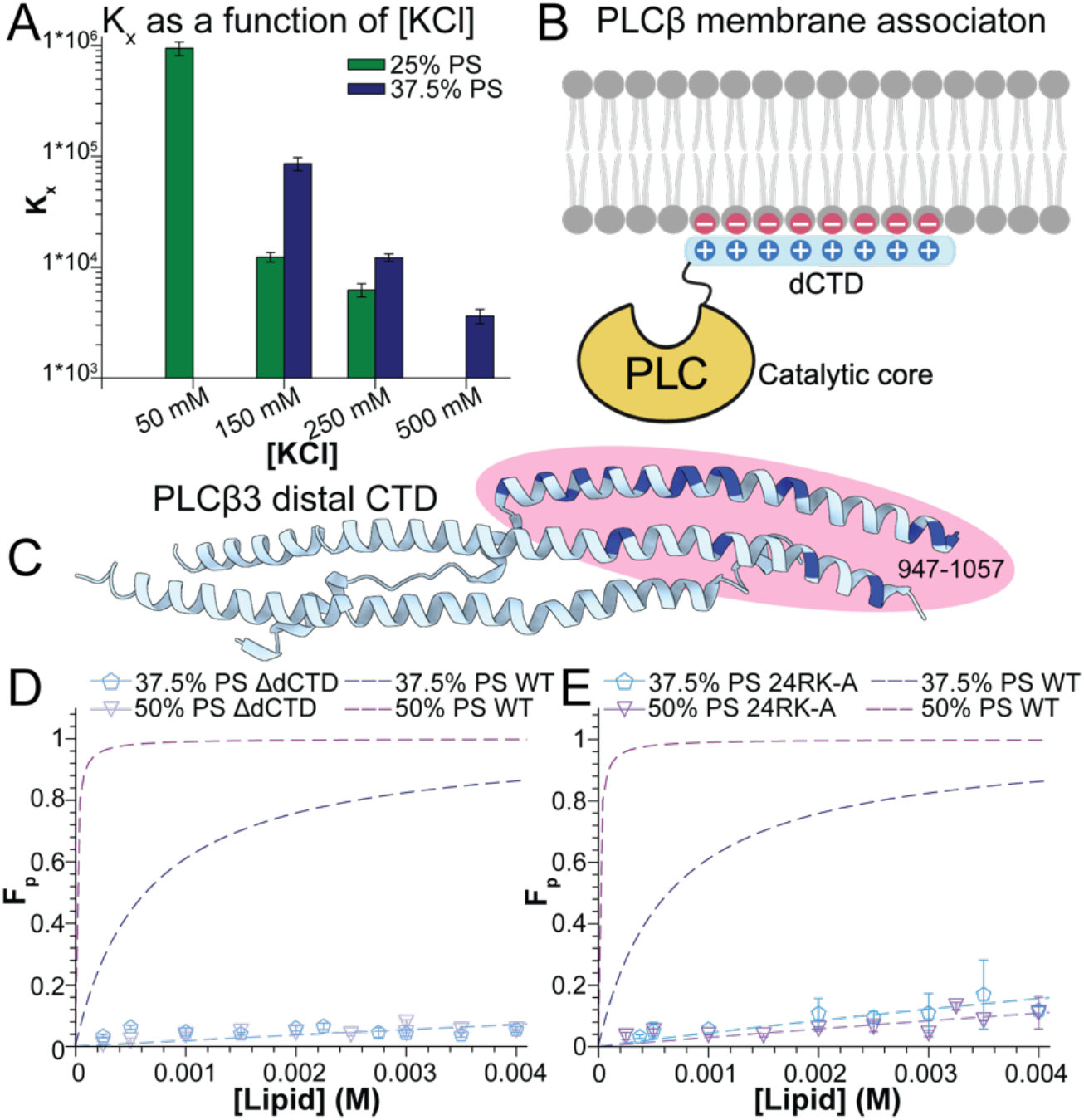
Arginine and lysine residues in the PLCβ dCTD mediate electrostatic interactions with the membrane. **A**: Comparison of K_x_ as a function of salt concentration for 50% PS (purple) or 37.5% PS (blue). Error bars are the error from the fit to determine K_x_. **B**: Cartoon representation of electrostatic interactions between basic residues in the dCTD and anionic lipids. **C**: PLCβ dCTD structure (PDB: 4GNK) with the 24 arginine and lysine residues within the region critical for membrane binding colored dark blue. **D-E**: Membrane partitioning curves for ΔdCTD (D) or 24RK-A (E) PLCβ in the presence of 37.5% PS or 50% PS. Symbols are individual lipid concentrations across at least 2 PLCβ concentrations and dashed lines are fits to Equation 2 with values reported in Table 1. Error bars are range of mean. The fit for PLCβ WT for 37.5% PS and 50% PS (from Fig. 1D) are shown for reference.

Numerous previous studies have shown that the dCTD of PLCβ is essential for membrane association and our previous structural studies have shown a direct interaction between this domain and the membrane (Schnabel et al., 1993; Wu et al., 1993; Kim et al., 1996; Adjobo-Hermans et al., 2013; Lyon and Tesmer, 2013; Falzone and MacKinnon, 2023a) (**Fig. 3B-C**). This domain also harbors numerous basic residues, which likely mediate the electrostatic interactions with anionic lipids. To investigate the role of this domain on the dependence of anionic lipids, we removed it (truncation after position 890) generating a ΔdCTD construct and assessed the membrane association in high proportions of anionic lipids (**Fig. S4A**). As predicted by our hypothesis, K_x_ for the truncated construct was significantly reduced, even in the presence of 50% anionic lipids (~82-fold in 37.5% PS to ~5000-fold in 50% PS) (**Fig. 3D, S4C-E, Table 1**). These results suggest that there are no additional electrostatic interactions between non-substrate anionic lipids and the catalytic core that can mediate robust membrane recruitment.

The dCTD of PLCβ3 harbors 56 basic residues and we set out to identify the critical ones, initially focusing on amino acids 947-1057, which was previously shown to be the minimal region required for membrane association (**Fig. 3B-C**) (Wu et al., 1993). The 24 basic residues in this stretch mostly line up on the same side of the helical bundle, consistent with this region serving as the primary membrane binding surface (**Fig. 3C**). We mutated all 24 basic residues in this stretch to alanine (24RK-A PLCβ) and assessed the partitioning. Even in the presence of 50% PS, partitioning was reduced ~2000-fold, comparable to the ΔdCTD construct (**Fig. 3E S4C, F-G, Table 1**), indicating that these basic residues mediate the critical electrostatic interactions between PLCβ and the membrane that underlie membrane recruitment.

### Anionic lipids and Gβγ cooperate to regulate PLCβ localization and function

Given that both anionic lipids and Gβγ contribute to PLCβ membrane association, we next addressed whether these two recruiters cooperate to enhance membrane localization. We reconstituted Gβγ into vesicles with increasing PS content including 100% PC, 10% PS, and 25% PS (**Fig. 4, S5**). In agreement with our hypothesis, the extent of Gβγ-dependent membrane recruitment was directly dependent on the proportion of anionic lipids, with low recruitment enhancement in 100% PC and 10% PS (~6-fold and ~3-fold respectively), and more substantial recruitment (~20-fold) in the presence of 25% PS (**Fig. 4, S5, Table 1**). These observations are consistent with previous results where we observed robust Gβγ-induced membrane recruitment in the context of the mixed lipid composition 2PE:1PC:1PS (Falzone and MacKinnon, 2023a). These studies demonstrate a functional crosstalk between G proteins and anionic lipids to regulate PLCβ localization.

**Figure 4:**
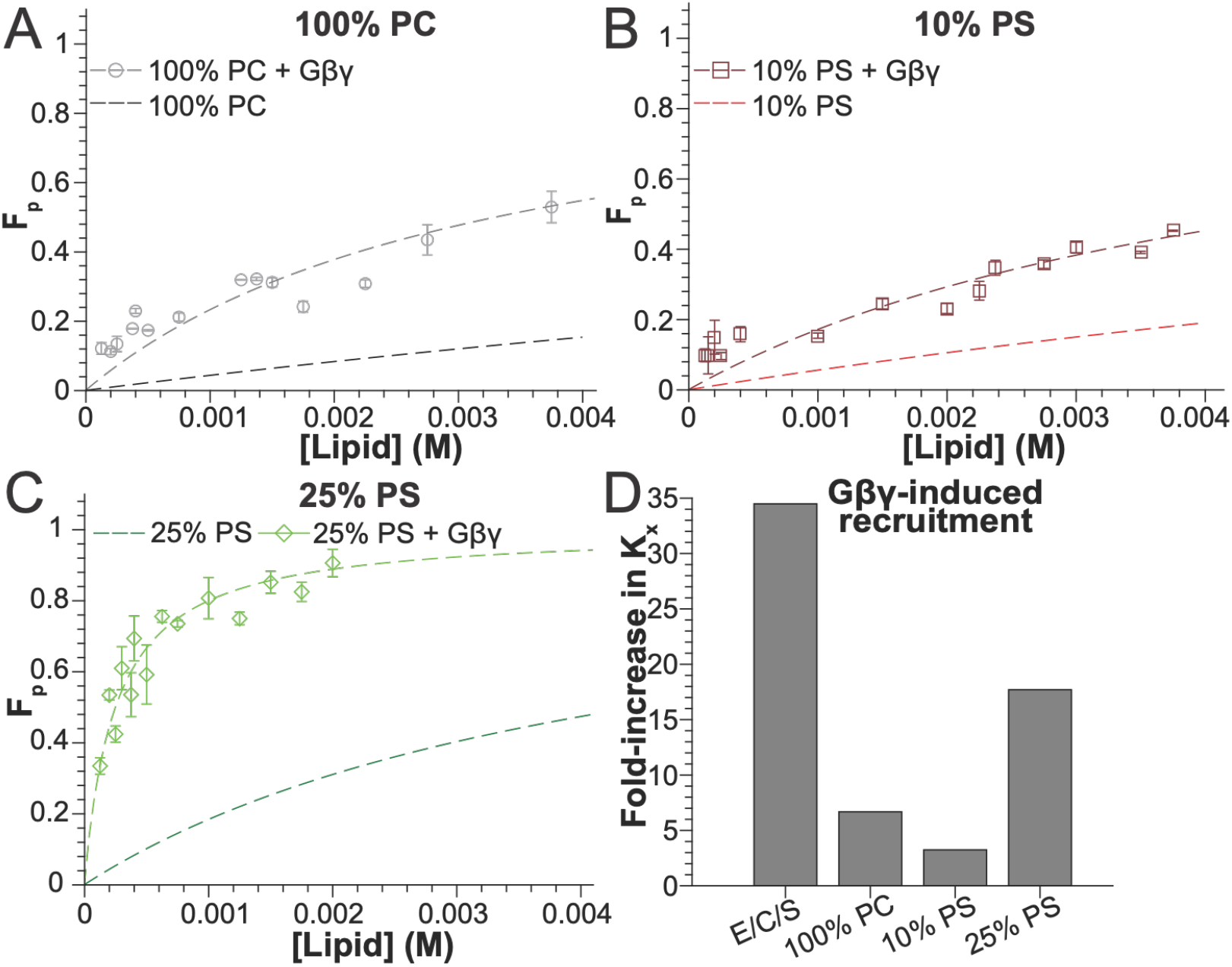
Gβγ and anionic lipids cooperate to fine-tune PLCβ membrane association. **A-C**: Membrane partitioning curves for PLCβ in the presence of 100% PC (A), 10% PS (B), or 25% PS (C) reconstituted with Gβγ. The fit for the respective condition in the absence of Gβγ is shown for reference. Symbols are individual lipid concentrations across at least 2 PLCβ concentrations and dashed lines are fits to Equation 2 with values reported in Table 1. Error bars are range of mean. **D**: Comparison of the Gβγ-induced fold-increase in membrane association across different lipid compositions. The previously-reported fold-increase in 2PE:1PC:1PS lipids is shown for comparison (Falzone and MacKinnon, 2023a).

## Discussion

Given the importance of membrane association for PLCβ function and regulation, our goal was to define the contributions of PLCβ interactions with non-substrate lipids, which is currently poorly understood (Romoser et al., 1996; Runnels et al., 1996; Jenco et al., 1997). Using our quantitative membrane partitioning assay, we demonstrate that anionic lipids are specifically required for robust membrane association, where polar lipids (PE) cannot substitute (**Fig. 1, S1, Table 1**). Further, PLCβ membrane association has a steep dependence on the concentration of anionic lipids, with a ~200 to ~400-fold increase in partitioning with only a 2-fold increase in PS or PG concentration (**Fig. 1D-E, Fig. 2F Table 1**). 50% anionic lipids is the highest concentration we can reliably measure, but this is likely not the saturation point, which prevents us from determining an EC_50_ value. However, the data presented suggest that the relationship between membrane association and anionic lipid content is highly cooperative, consistent with the idea of multiple contacts between the protein and the membrane.

Further, we demonstrate that this dependence on anionic lipids is mediated by the N-terminal portion of the dCTD, specifically basic residues between positions 947-1057 in PLCβ3 (**Fig. 3C-E S4C-G**), consistent with previous results highlighting the importance of this region for membrane association (Schnabel et al., 1993; Wu et al., 1993; Kim et al., 1996; Adjobo-Hermans et al., 2013; Lyon and Tesmer, 2013). PLCβ constructs lacking the dCTD retain catalytic activity (Lyon and Tesmer, 2013), suggesting that the catalytic core itself can interact with the membrane to some degree. However, our results suggest that interactions between the catalytic core and the membrane are not mediated by anionic non-substrate lipids, given that 50% PS content could not rescue partitioning of constructs with missing or mutated dCTDs. Accordingly, it is likely that these interactions are mediated by PIP2 in the active site, but further experiments are required to evaluate this proposal.

Finally, we demonstrate the anionic lipids and Gβγ cooperate to enhance PLCβ membrane association, where the magnitude of Gβγ-dependent recruitment observed is dependent on the content of anionic lipids, with low recruitment (~6-fold and ~3-fold) observed with low anionic lipids and ~20-fold observed with 25% anionic lipids (**Fig. 4, S5, Table 1**). These results are consistent with our previous proposal that membrane-associated Gβγ shifts the equilibrium of PLCβ to more membrane-associated (Falzone and MacKinnon, 2023a; Falzone et al., 2026). We previously demonstrated robust Gβγ-dependent membrane recruitment in reconstituted systems (Falzone and MacKinnon, 2023b) and in the native cellular context (Falzone et al., 2026). However, these observations are contrary to several other studies where this recruitment is not observed, which has left the topic of Gβγ-dependent membrane recruitment controversial (Romoser et al., 1996; Jenco et al., 1997; Senarath et al., 2025; Fisher et al., 2026). Additionally, our previous structural studies demonstrate that Gβγ dramatically re-orients the PLCβ catalytic core on the membrane surface, making it more poised for catalysis, which is undoubtably an important component of its activation mechanism (Falzone and MacKinnon, 2023a). Based on observations in this work, we propose an updated framework for the molecular mechanisms underlying PLCβ regulation by Gβγ, where both recruitment and reorientation of the catalytic core play important roles depending on the cellular conditions (**Fig. 5**). When PLCβ membrane association is low, i.e., in the context of low anionic lipids, Gβγ is unable to provide a major shift in the PLCβ equilibrium between the solution and the membrane, and it thus facilitates activation via orienting the catalytic core for catalysis. When PLCβ membrane association is intermediate, i.e., in the context of high anionic lipid content, sufficient PLCβ-Gβγ complex can shift the PLCβ equilibrium to membrane associated, thereby facilitating activation by both membrane recruitment and catalytic core orientation. In support of this proposal, in cases where Gβγ-dependent recruitment is not observed, only modest increases in catalytic activity (~3-fold) are observed (Romoser et al., 1996; Runnels et al., 1996; Jenco et al., 1997; Fisher et al., 2020), whereas in cases with clear Gβγ-dependent recruitment, hydrolysis stimulation is much more substantial (~20-70-fold) (Falzone and MacKinnon, 2023a), consistent with singular vs dual activation mechanisms. Thus, orientation of the catalytic core on the membrane surface is likely the major activation mechanism in cases where Gβγ-dependent membrane recruitment is not observed under conditions where it stimulates PIP2 hydrolysis. Accordingly, in light of the large differences in PLCβ membrane association observed in this work in the context of different salt concentrations as well as different lipid compositions and concentrations, we propose that the reported disagreements are due to differences in experimental conditions and we hope this work provides some clarity to these discrepancies (Romoser et al., 1996; Runnels et al., 1996; Jenco et al., 1997; Campbell et al., 2003; Fisher et al., 2020).

**Figure 5:**
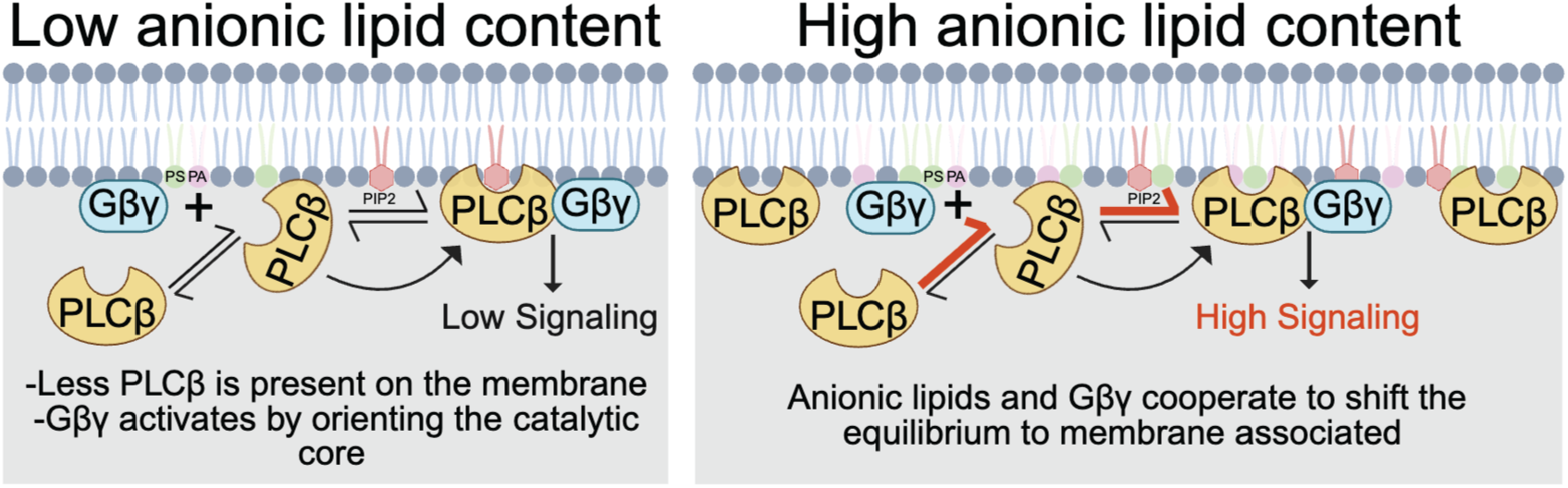
Model for context-dependent regulation of PLCβ enzymes by Gβγ. In cases of low PLCβ membrane association, i.e., in the presence of low anionic lipid content (left), Gβγ is unable to promote a significant shift in the PLCβ membrane-solution equilibrium and activation is facilitated by orientation of the catalytic core on the membrane surface. In cases of moderate PLCβ membrane association, i.e., in the presence of intermediate anionic lipid content, sufficient PLCβ-Gβγ complex forms and is able to shift the PLCβ equilibrium to membrane associated, as well as orient the catalytic core for catalysis.

In cells, anionic lipids comprise ~20% of the inner leaflet of the plasma membrane in total, but the plasma membrane is highly organized, likely harboring locally higher and lower concentrations (Platre and Jaillais, 2017; Robinson et al., 2019; Levental and Lyman, 2023). Further, anionic lipid content is under the control of a myriad of signaling pathways, including RTK and GPCR signaling, which regulate lipase enzymes (Platre and Jaillais, 2017; Robinson et al., 2019; Levental and Lyman, 2023). Given the steep dependence of membrane association on anionic lipids between 25% and 50%, this system is poised to be regulated by signaling-dependent local changes in anionic lipid content. A prime example is PLD enzymes, which cleave PC to produce PA, a critical mediator of cellular signaling as well as anionic lipid content (Platre and Jaillais, 2017; Robinson et al., 2019; Levental and Lyman, 2023). Finally, cytoplasmic-facing intracellular membranes, such as the ER and Golgi, have a much lower anionic lipid content (<10%) (Bigay and Antonny, 2012; Levental and Lyman, 2023). Thus, the requirement of anionic lipids for robust PLCβ membrane association could be an additional mechanism to prevent targeting of these enzymes to internal membranes, which comprise the majority of lipids in the cell, where their substrate is generally not accessible. More broadly, our findings establish membrane electrostatics as a fundamental regulator of PLCβ signaling, revealing how changes in membrane composition can be integrated with receptor-driven pathways to dynamically regulate cellular signal transduction.

## Materials and Methods

### Constructs and Mutagenesis

The human PLCβ3 gene was provided by Dr. Sondek (Charpentier et al., 2014) and contained residues 10 to 1234. The PLCβ3 ΔdCTD construct was generated by truncation via the KLD enzyme mix from NEB. The PLCβ3 24RK-A was synthesized by twist bioscience and cloned into a pfastbac vector with an N-terminal GFP tag (identical to the wildtype PLCβ3) (Falzone and MacKinnon, 2023b; Falzone and MacKinnon, 2023a) using Gibson assembly. Mutated positions include: R958A, K961A, K965A, R967A, R969A, R972A, R975A, R978A, K979A, K980A, K983A, K984A, R990A, R991A, R1004A, R1006A, R1008A, K1022A, K1029A, R1030A, R1037A, R1046A, R1056A, and R1057A. All sequences were confirmed and assessed for off-target effects using whole-plasmid sequencing.

### Preparation of GFP nanobody-coupled Sepharose resin

The enhancer GFP nanobody (Kirchhofer et al., 2010) with N-terminal Glutathione S-transferase (GST) and His tags in a pGEX-6P-1 backbone (pHN922) was a gift from Kazuhiro Abe (Addgene plasmid # 178975 ; http://n2t.net/addgene:178975; RRID:Addgene_178975). The plasmid was transformed into BL21 (DE3) competent cells according to the manufacturers protocol. Colonies were used to initiate a starter culture in 2xYT media supplemented with 0.4% glucose and 100 µg/mL ampicillin grown at 30°C overnight. The starter culture was used to inoculate large cultures of 2xYT media supplemented with 0.4% glucose and 100 µg/mL ampicillin, which were grown to an OD600 of 0.6 at 30°C. Expression was induced with 0.15 mM isopropyl-β-D-thiogalactoside (IPTG) and the temperature was reduced to 20°C for ~24 hours. Cells were harvested by centrifugation at 3,500 x *g* for 10 minutes. Pellets were flash frozen and stored at -80°C until use. Purification was carried out at 4°C. Cells were resuspended in PBS (10 mM Na-phosphate pH 7.5, 137 mM NaCl, and 2.7 mM KCl. pH 7.4), supplemented with 15 mM imidazole, DNase, and protease inhibitors (12.5 μg/mL leupeptin, 12.5 μg/mL pepstatin A, 625 μg/mL AEBSF, 1 mM benzamidine, 100 μg/mL trypsin inhibitor, 1x aprotinin, and 1 mM PMSF) and lysed by brief sonication. Lysate was clarified by centrifugation at 39,000 x *g* for 45 minutes and bound in batch to 10 mL NiNTA resin equilibrated with PBS supplemented with 15 mM imidazole for one hour. The resin was washed in batch with 100 mL of equilibration buffer then loaded onto a column and washed by gravity flow with 10 column volumes of PBS supplemented with 30 mM imidazole. Protein was eluted with ~70 mL of PBS supplemented with 250 mM imidazole. Protein was concentrated to 30 mL using a 15-mL Amicon concentrator with 10-kDa molecular weight cutoff and buffer exchanged using PD-10 desalting columns into PBS to remove the imidazole, flash frozen, and stored at -80°C until coupling.

Protein was coupled to CNBr-Activated Sepharose 4 Fast Flow resin according to the manufacturer’s protocol and as previously described (Falzone and MacKinnon, 2023b; Falzone and MacKinnon, 2023a). Briefly, 300 mg of protein was coupled to 25 g of resin, which resulted in ~200 mL of 50% slurry. Resin was prepared by mixing with 1 mM cold HCl, manually breaking up chunks, loaded onto a column and washed with additional cold HCl. The resin was then washed with cold coupling buffer (100 mM NaHCO_3_, and 500 mM NaCl) and transferred to a bottle. The nanobody was diluted into coupling buffer, added to the resin, and rotated overnight at 4°C. The resin was pelleted by spinning for 10 minutes at 2,000 *g* and washed with additional coupling buffer. The discarded supernatant was checked for uncoupled protein, and the resin was spun again, resuspended in blocking buffer (50 mM NaPhosphate pH 8.0, 150 mM NaCl, 50 mM glycine), and rotated at room temperature for 2 hours. The resin was washed extensively with wash buffer (10 mM NaPhosphate pH 7.0 and 150 mM NaCl) and finally stored at 4°C as a 50% slurry in wash buffer.

### PLCβ3 purification

All PLCβ constructs were expressed, purified, and fluorescently labeled as previously described (Falzone and MacKinnon, 2023b; Falzone and MacKinnon, 2023a). Briefly, PLCβ3 genes in the pFastBac vector with an N-terminal GFP tag were transformed into DH10bac cells to produce bacmid DNA, which was used to transfect Sf9 cells to produce baculovirus. High Five insect cells at 2×10^6^ cells/mL were infected with 15-25 mL of P3 virus per liter of culture and harvested 36-48 hours after infection by centrifugation at 3,500 x *g* for 15 minutes. Pellets were flash frozen and stored at -80°C until use. Purification was carried out at 4°C. For functional experiments with non-labeled PLCβ3, cells were resuspended in buffer containing 50 mM HEPES pH 8.0, 50 mM NaCl, 10 mM 2-mercaptoethanol, 5% glycerol (v/v), 0.1 mM EDTA, 0.1 mM EGTA, DNase and protease inhibitors (12.5 μg/mL leupeptin, 12.5 μg/mL pepstatin A, 625 μg/mL AEBSF, 1 mM benzamidine, 100 μg/mL trypsin inhibitor, 1x aprotinin, and 1 mM PMSF) and lysed by brief sonication. Lysate was clarified by centrifugation at 39,000 x *g* for 45 minutes and bound to GFP nanobody-coupled Sepharose resin (prepared in-house) for one hour. The resin was washed in batch once with 10 column volumes of buffer containing 20 mM HEPES pH 8.0, 400 mM NaCl, 10 mM 2-mercaptoethanol, 2% glycerol (v/v), 0.1 mM EDTA, 0.1 mM EGTA, and protease inhibitors (625 μg/mL AEBSF, 1 mM benzamidine, 100 μg/mL trypsin inhibitor, and 1x aprotinin) then loaded onto a column and washed with an additional 10 column volumes by gravity flow. Protein was eluted by cleavage with 3C PreScission protease (made in house) for two hours, concentrated to ~1 mL using a 15-mL Amicon concentrator with 100-kDa molecular weight cutoff, and further purified by size exclusion chromatography using a Superdex 200 10/300 increase column equilibrated with buffer containing 20 mM HEPES pH 8.0, 100 mM NaCl, 5 mM Dithiothreitol (DTT), 2% glycerol (v/v), 0.1 mM EDTA, 0.1 mM EGTA, and protease inhibitors (12.5 μg/mL leupeptin, 12.5 μg/mL pepstatin A, 625 μg/mL AEBSF, 1 mM benzamidine, 100 μg/mL trypsin inhibitor, 1x aprotinin, and 1 mM PMSF). Fractions with PLCβ3 were pooled, flash frozen, and stored at -80°C for later use.

To purify protein for non-specific cysteine labeling, the 10 mM 2-mercaptoethanol in all buffers was replaced with 2 mM tris(2-carboxyethyl)phosphine (TCEP) and the buffer for labeling was prepared with miliQ water treated with Chelex 100 sodium form (1 g Chelex per 100 mL H_2_O for 1 hour). Following elution, the protein was buffer exchanged using PD-10 desalting columns into Chelex-treated labeling buffer containing 20 mM HEPES pH 7.4, 400 mM NaCl, 2 mM TCEP, 2% glycerol (v/v), 0.1 mM EDTA, 0.1 mM EGTA, and protease inhibitors (625 μg/mL AEBSF, 1 mM benzamidine). Sulfo-Cyanine5 maleimide (Lumiprobe) was added in 5-fold molar excess and incubated overnight protected from light. Labeled protein was concentrated to ~1 mL using a 15-mL Amicon concentrator with 100-kDa molecular weight cutoff, and further purified by size exclusion chromatography using a Superdex 200 10/300 increase column in buffer containing 20 mM HEPES pH 8.0, 100 mM NaCl, 5 mM Dithiothreitol (DTT), 2% glycerol (v/v), 0.1 mM EDTA, 0.1 mM EGTA, and protease inhibitors (12.5 μg/mL leupeptin, 12.5 μg/mL pepstatin A, 625 μg/mL AEBSF, 1 mM benzamidine, 100 μg/mL trypsin inhibitor, and 1x aprotinin). Fractions with labeled PLCβ3 were pooled, flash frozen, and stored at -80°C for later use. Labeling efficiency was consistently 40-100%.

### Gβγ purification

Untagged human Gβ1 was co-expressed with bovine Gγ2 with an N-terminal His-YFP tag (or His-tag alone) in High Five insect cells using 12 and 8 mL of P3 baculovirus respectively at 2×10^6^ cells/mL for 36-48 hours as previously described (Falzone and MacKinnon, 2023b; Falzone and MacKinnon, 2023a). Briefly, cells were harvested by centrifugation at 3,500 x *g* for 15 minutes and pellets were flash frozen and stored at -80°C until use. Purification was carried out at 4°C. Cells were resuspended in buffer containing 25 mM Tris-HCl pH 8.0, 125 mM NaCl, 5 mM EGTA, 5 mM DTT, DNase, and protease inhibitors (12.5 μg/mL leupeptin, 12.5 μg/mL pepstatin A, 625 μg/mL AEBSF, 1 mM benzamidine, 100 μg/mL trypsin inhibitor, 1x aprotinin, and 1 mM PMSF) and broken by manual homogenization. Membranes were separated by centrifuging at 39,000 x *g* for 30 minutes, resuspended in buffer containing 25 mM Tris-HCl pH 8.0, 125 mM NaCl, DNase and protease inhibitors (12.5 μg/mL leupeptin, 12.5 μg/mL pepstatin A, 625 μg/mL AEBSF, 1 mM benzamidine, 100 μg/mL trypsin inhibitor, 1x aprotinin, and 1 mM PMSF) and manually homogenized again. Proteins were extracted using 1% sodium cholate added from a 10% stock by mixing for one hour. Lysate was clarified by centrifuging at 39,000 x *g* for 30 minutes and bound in batch to 10 mL TALON resin equilibrated with buffer containing 25 mM Tris-HCl pH 8.0, 125 mM NaCl, and 1% sodium cholate for one hour. The resin was washed in batch with 100 mL of equilibration buffer then loaded onto a column and washed by gravity flow with 10 mM imidazole buffer (25 mM Tris-HCl pH 8.0, 125 mM NaCl, 1% sodium cholate, and 10 mM imidazole). Protein was eluted with ~20 mL buffer containing 25 mM Tris-HCl pH 8.0, 125 mM NaCl, 1% sodium cholate, and 200 mM imidazole and concentrated to ~2 mL using a 15-mL Amicon concentrator with 30-kDa molecular weight cutoff. The final 2 mL of protein was diluted to ~20 mL using buffer without imidazole and incubated with 3C PreScission protease (prepared in-house) overnight at 4°C remove the His-YFP. The cleaved protein was passed over a column of 10 mL TALON resin equilibrated with 25 mM Tris-HCl pH 8.0, 125 mM NaCl, 1% sodium cholate, and 20 mM imidazole three times to remove the free His-YFP and concentrated to 1 mL using a 15-mL Amicon concentrator with 30-kDa molecular weight cutoff. Gβγ was purified further via size exclusion chromatography using a Superdex 200 10/300 increase column in buffer containing 25 mM Tris-HCl pH 8.0, 125 mM NaCl, 1% sodium cholate, and 5 mM DTT. Fractions containing Gβγ were pooled and concentrated to 5-10 mg/mL using 4-mL Amicon concentrator with 30-kDa molecular weight cutoff and immediately used for reconstitution.

### Liposome Reconstitution

Reconstitutions for partitioning included 0.1% (wt/wt) of 1,2-dioleoyl-sn-glycero-3-phosphoethanolamine-N-(lissamine rhodamine B sulfonyl) (18:1 Liss Rhod PE). Lipid mixtures were calculated using mass. Lipids in chloroform were mixed according to the desired ratios and dried under a stream of argon, washed with pentane, dried under a stream of argon, and incubated under vacuum for at least one hour. Lipids were resuspended at 25 mM in buffer containing 25 mM HEPES pH 7.4, 150 mM KCl and 5 mM DTT and sonicated to clarity. 40 mM sodium cholate was added, and the mixture was sonicated briefly. Lipids were diluted to a final concentration of 12.5 mM, maintaining 40 mM cholate, prior to initiating detergent removal. For Gβγ-containing liposomes, Gβγ was added at a protein to lipid ratio of 1:5 (wt/wt) as the lipid concentration was reduced to 12.5 mM maintaining 40 mM sodium cholate and incubated at 4°C for one hour. Detergent was removed using four exchanges of 200 mg/mL biobeads washed with reconstitution buffer after two hours, 12 hours, two hours, and two hours at 4°C. Liposomes were flash frozen and stored at -80°C until use. For experiments with different salt concentrations, the salt was changed throughout the reconstitution and portioning experiments. Reconstitutions for functional assays were prepared using the same procedure, except the final lipid concentration was 7.5 mM upon detergent removal and lipids were supplements with 2 mol% PIP2 (18:1 PI(4,5)P2).

### NMR experiments to measure lipid concentration

A small volume of liposomes from reconstitutions (7 μL) was dissolved in ~250 μL of a mixture of deuterated methanol and chloroform (5:6) containing 100 μM of the standard sodium trimethylsilyl propionate (TSP). Proton spectra were measured on a Bruker 500 MHz instrument equipped with an AVANCE III console and a 5 mm TCI cryoprobe. Spectra were collected in 3 mm tubes at 298 K using a 30° flip angle, 16 scans, and 2.8 second acquisition time and a recycle delay of 18 μs. Spectra were processed using TopSpin 4.1.1 for line broadening, phasing, and baseline correction. The triplet lipid -CH_3_ peak at 0.875 ppm was integrated relative to the TSP peak at 0 ppm and normalized to the difference in protons (nine for TSP and 6 for the lipid CH3). The normalized peak area was used to determine the lipid concentration using the known 100 μM concentration of TSP as previously described (Falzone and MacKinnon, 2023a).

### PLCβ vesicle partitioning experiments

Partitioning experiments were carried out as described (Falzone and MacKinnon, 2023b; Falzone and MacKinnon, 2023a). Briefly, reconstituted liposomes with or without Gβγ were subjected to 10 freeze-and-thaw cycles and extruded 21 times through a 200 nm membrane to produce LUVs. Fixed concentrations of lipids were mixed with LD655-labeled PLCβ3 in buffer containing 25 mM HEPES pH 7.4, 150 mM KCl, and 5 mM DTT, incubated for one hour, and centrifuged for one hour at 100,000 x *g* at room temperature. The supernatant was removed and the membrane pellet was resuspended in an equal volume of buffer. The input, pellet, and supernatant samples were analyzed by SDS-PAGE and imaged using in-gel fluorescence to detect Cy5-labeled PLCβ3 (**Fig. S1-S5**). Additionally, input, supernatant, and pellet samples were solubilized in 5% Anapoe-C12E10 to eliminate scattering artifacts and the fluorescence signal was measured for Cy5 (Ext-637, Em-667) and Rhodamine (Ext-550, Em-583) using a BMG LABTECH plate reader. The Rhodamine signal was used to estimate the fraction of lipids that were pelleted and the measurements were corrected for this as well as the loss of material using the difference between the input and output (pellet and supernatant) Cy5 signal. The reported lipid concentration is 50% of the total lipid concentration added in solution because PLCβ3 only has access to the outer leaflet. Each lipid concentration was repeated with two different PLCβ3 concentrations, ranging from 50 nM to 250 nM. Values for fraction of PLCβ partitioned (F_p_), were determined, plotted against lipid concentration, and fit to Equation 2 to determine K_x_. All K_x_ values (Table 1) are from at least 7 different lipid concentrations across 2 different reconstitutions and partitioning measurements.

### DLS

For DLS measurements, lipids were prepared as for partitioning (subjected to 10 freeze-and-thaw cycles and extruded 21 times through a 200 nm membrane to produce LUVs). LUVs were diluted 1:25 in 100 µL of reconstitution buffer and measured using a DynaPro NanoStar dynamic light scattering instrument (Wyatt, CA) at 20°C with a laser power of 2% using a low-volume quartz cuvette. For each sample, 22 10-second acquisitions were collected and averaged. Data were analyzed using the algorithms available in the instrument control software, using the “no molecular weight” mode, and exported as size distributions and average sizes. Three independent reconstitutions and preparations were measured for each lipid composition.

### PLCβ lipase activity measurements

For lipase assays using XY-69, lipids were prepared as for partitioning (subjected to 10 freeze-and-thaw cycles and extruded 21 times through a 200 nm membrane to produce LUVs). XY-69 was added to LUVs at 0.1 mol%. Lipids were diluted to a final concentration of 500 µM with reconstitution buffer (25 mM HEPES pH 7.4, 150 mM KCl) supplemented with 100 µM CaCl_2_ and distributed into black 384-well plates. The XY-69 signal was measured using a BMG LABTECH plate reader (Ext-485, Em-520) over time. The baseline signal was measured for ~30s and then PLCβ was manually added and mixed in each well at a final concentration of 10 nM and the measurement was continued. Each experiment consists of three identical wells, which were averaged and treated as technical replicates. Three technical replicates on three separate experimental days were completed for each lipid composition. The 50% PS and 25% PS, the signal was well-fit by a standard double exponential function and fitting was carried out accordingly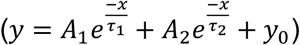. The curves from 100% PC could not be fit by a double exponential function and instead were fit by a combined exponential and linear function 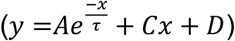 where C is the slope of the linear component and D is the offset. Parameters from the fits are shown in **Fig. 2C, and S1G-I**.

## Data Availability

Data are available in the article itself and its supplementary materials.

## Acknowledgements

We thank all members of the Falzone lab for helpful discussions. We thank Philipp Schmideter for helpful discussions and feedback on the manuscript. We thank Reuben Harris and members of his lab (Allen York and Agnieszka Dabrowska) for use of their cell culture facility. We thank Patrick Sung and members of his lab (Stephen Holloway) for supplies and support over the course of the project. We thank David Libich and members of his lab (Xiaoping Xu and Hairan Weng) for use of the dynamic light scattering system. NMR experiments were conducted in the Structural Biology Core Facilities, a part of the Institutional Research Cores at the University of Texas Health Science Center at San Antonio supported by the Office of the Vice President for Research and the Mays Cancer Center Drug Discovery and Structural Biology Shared Resource (NIH P30 CA054174 and NIH P01CA275717) and by NIH Award Number S15OD039797. T.G.G. is a trainee of the South Texas Undergraduate Research Opportunities Program (STUROP) at UT Health San Antonio and St. Mary’s University. This work is supported by the NIGMS T32GM145432 (South Texas Medical Scientist Training Program), a Research Grant from The Robert A. Welch Foundation: AQ-2233-20250403 (M.E.F), a CPRIT Recruitment of First Time Tenure Track Faculty Award: RR240042 (M.E.F), and The Voelcker Foundation Young Investigator Award (M.E.F).

## Author Contributions

M.E.F. and M.A.C. designed experiments, M.E.F., M.A.C., M.B.B., K.E.C., Y.Q., T.G.G., and R.M., performed the experiments and analyzed the data. M.E.F. and M.A.C. prepared the manuscript with input from all authors.

