## Supplementary material for "Anionic lipids regulate PLCβ membrane recruitment": Figure S

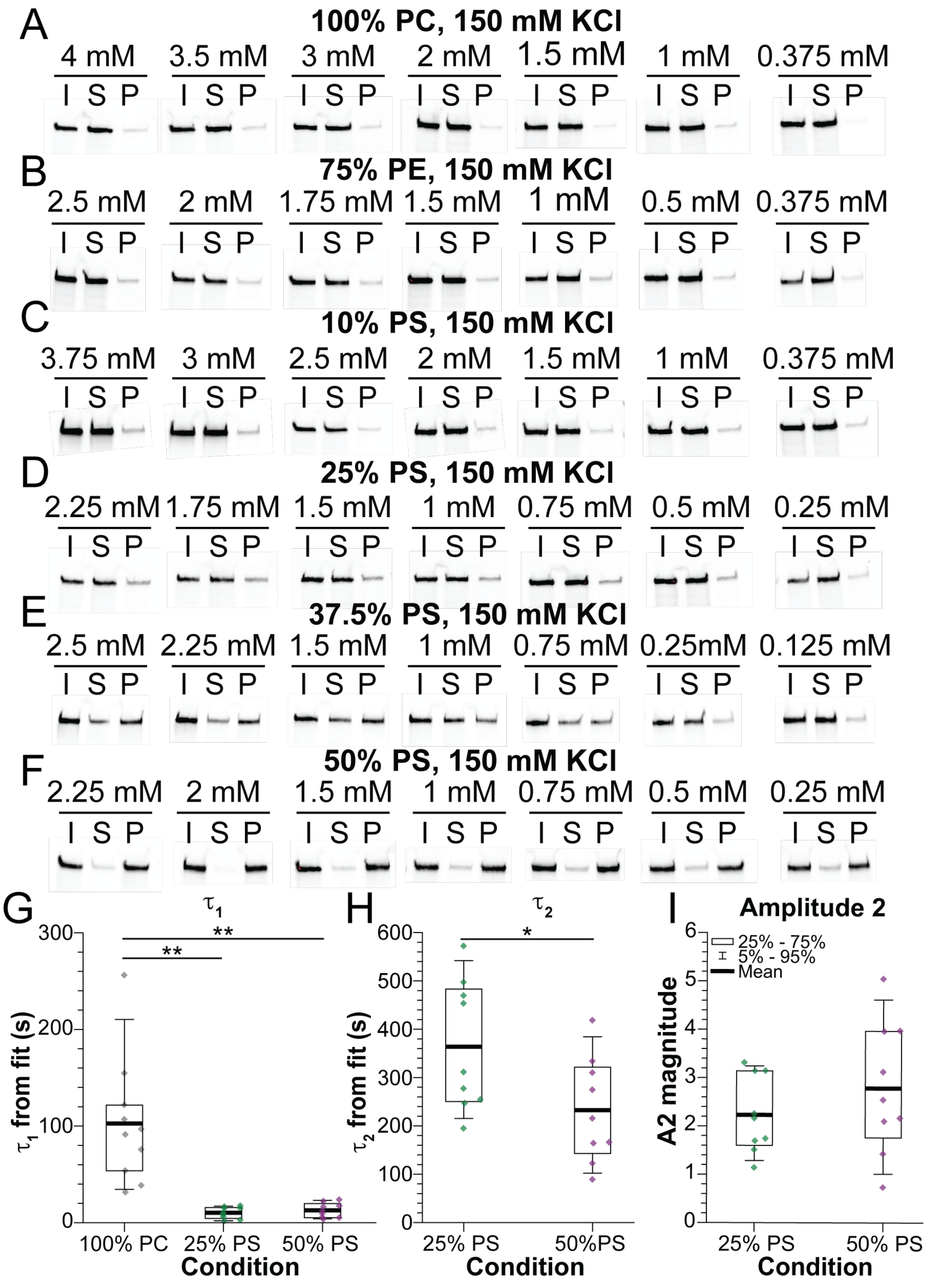


**Figure S1**: *PS significantly enhances PLCβ membrane partitioning*. **A-F**: Representative SDS-PAGE gels imaged for Cy5 fluorescence from partitioning experiments with 100% PC lipids (A), 75% PE (B), 10% PS (C), 25% PS (D), 37.5% PS (E), and 50% PS (F). I represents input, S represents supernatant, and P represents pellet. Final PLCβ concentration used in the experiments is 150 nM (A-D, F) or 250 nM (E). **G-I**: Comparison of fit parameters $\tau_1$ (G), $\tau_2$, and Amplitude 2 (I) from the XY-69 fluorescence experiments. Diamonds represent individual replicates. *: $0.05>p>0.005$. **: $0.005>p>0.0005$. ***: $p<0.0005$.

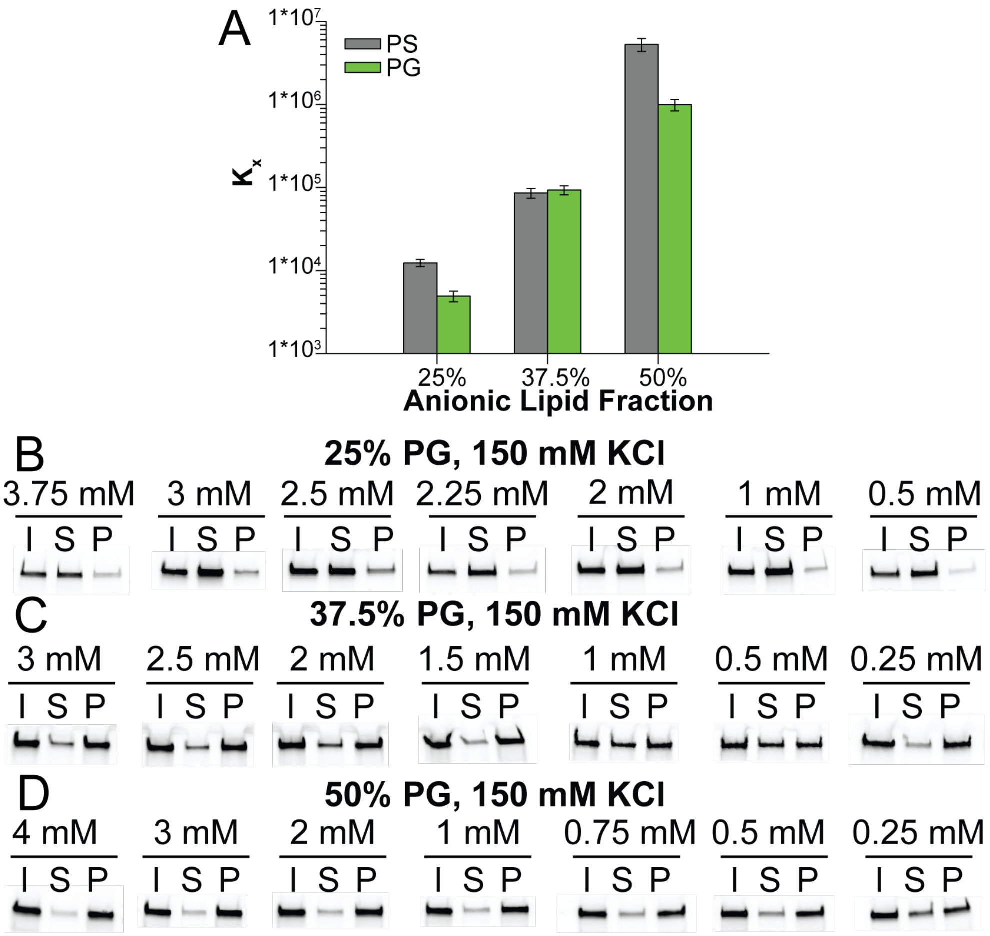


**Figure S2**: *PG lipids also support robust PLCβ partitioning*. **A**: Comparison of $K_x$ as a function of anionic lipid content across PS (gray) and PG (green). Error bars are the error from the fit to determine $K_x$. **B-D**: Representative SDS-PAGE gels imaged for Cy5 fluorescence from partitioning experiments with 25% PG lipids (B), 37.5% PG (C), and 50% PG (D). I represents input, S represents supernatant, and P represents pellet. Final PLCβ concentration used in the experiments is 150 nM.

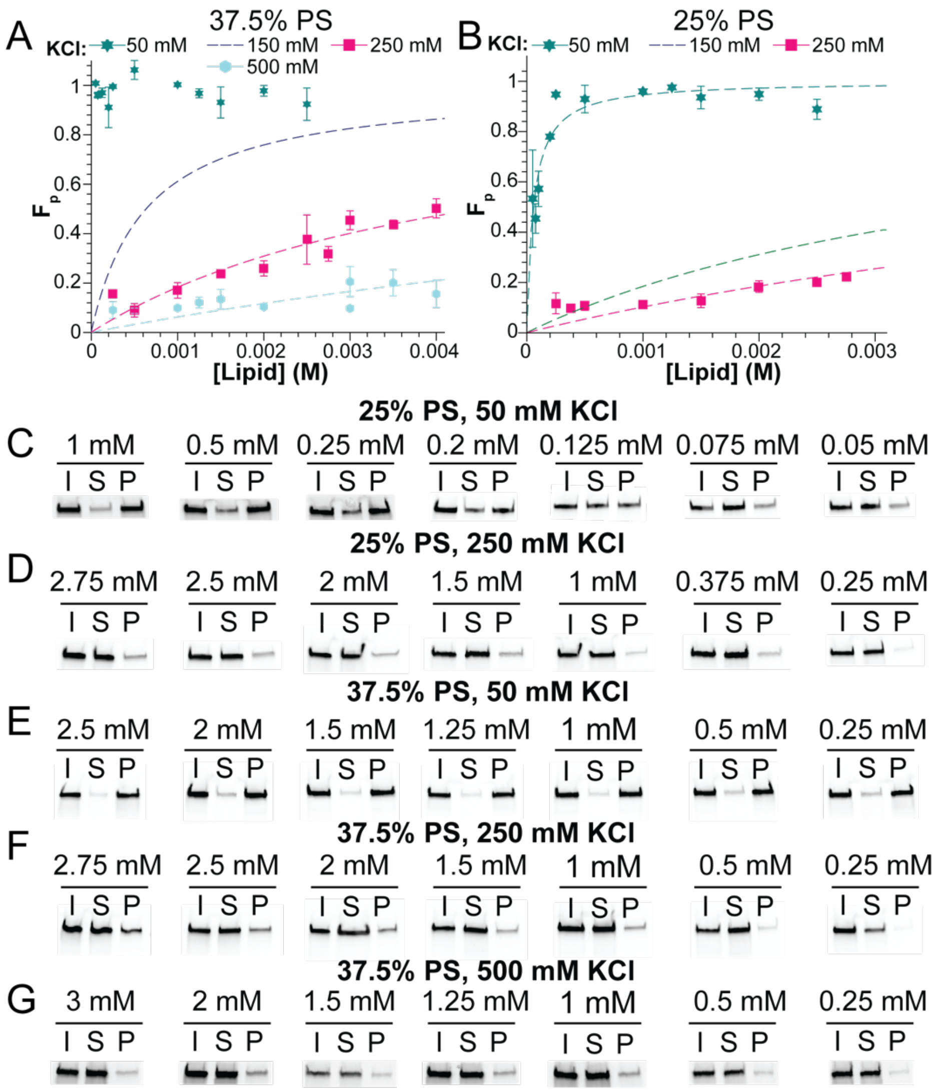


**Figure S3**: *PLCβ membrane partitioning is sensitive to salt concentration*. **A-B**: Membrane partitioning curves for PLCβ for LUVs comprised of 37.5% PS (A) or 25% PS (B) in 50 KCl mM (dark cyan), 250 mM KCl (pink), or 500 mM KCl (light blue). The fits for 37.5% PS and 25% PS in 150 mM KCl (from Fig. 1D) are shown for reference. Symbols are individual lipid concentrations across at least 2 PLCβ concentrations and dashed lines are fits to Equation 2 with values reported in Table 1. Error bars are range of mean. **C-G**: Representative SDS-PAGE gels imaged for Cy5 fluorescence from partitioning experiments with 25% PS lipids in 50 mM KCl (C), 25% PS lipids in 250 mM KCl (D), 37.5% PS lipids in 50 mM KCl (E), 37.5% PS lipids in 250 mM KCl (F), and 37.5% PS lipids in 500 mM KCl (G). I represents input, S represents supernatant, and P represents pellet. Final PLCβ concentration used in the experiments is 150 nM (C-D) or 50 nM (F-G).

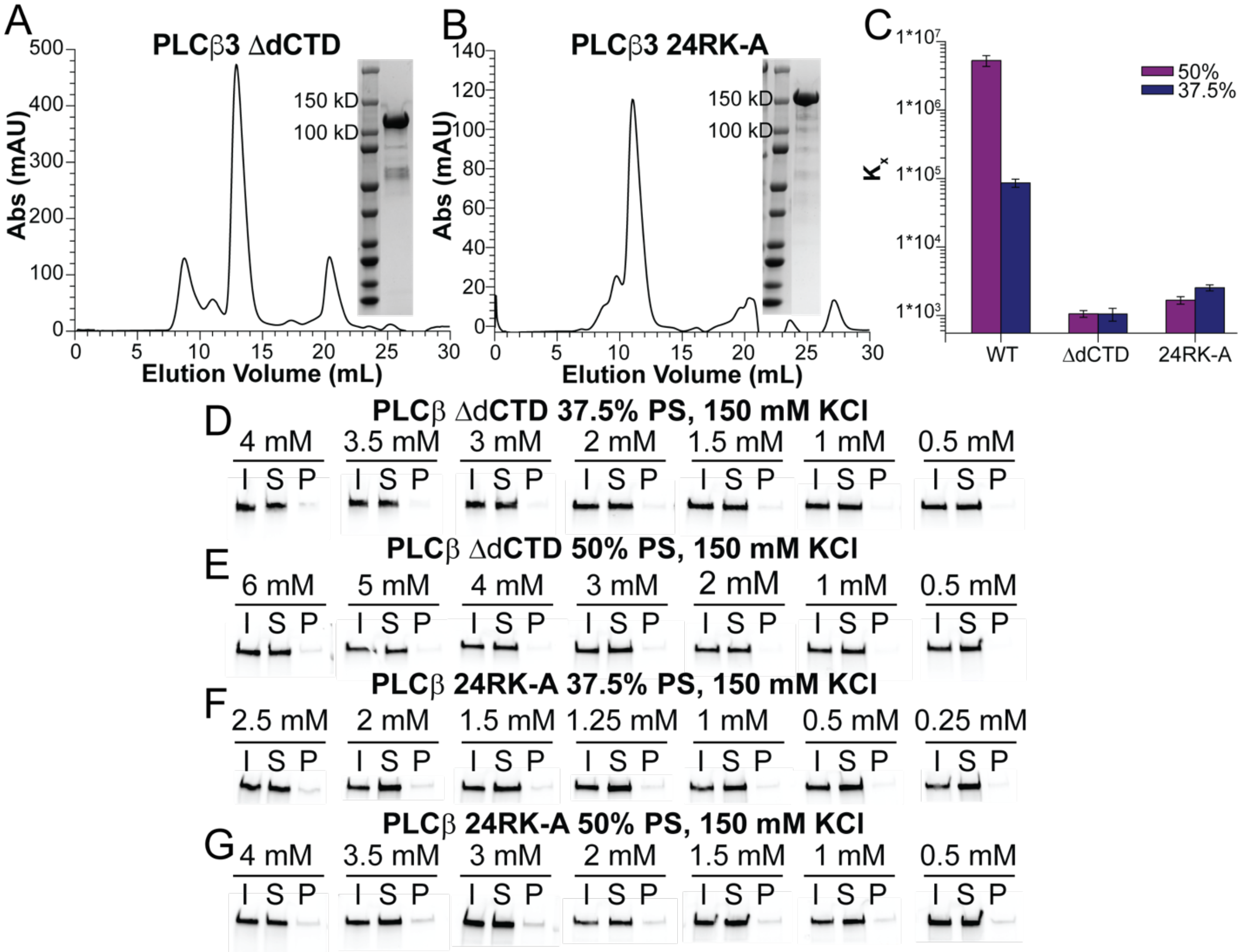


**Figure S4**: *The PLCβ3 dCTD mediates electrostatic interactions with anionic lipids*. **A-B**: Size exclusion chromatography profiles for the purification of PLCβ3 ΔdCTD (A) or PLCβ3 24RK-A using a Superdex 200 10/300 column. Inset shows SDS-PAGE gel of purified protein. **C**: Comparison of $K_x$ across wildtype, ΔdCTD, and 24RK-A PLCβ3 in LUVs with 50% PS (purple) or 37.5% PS (dark blue). Error bars are the error from the fit to determine $K_x$. **D-G**: Representative SDS-PAGE gels imaged for Cy5 fluorescence from partitioning experiments with PLCβ3 ΔdCTD (D-E) or PLCβ3 24RK-A (F-G) in 37.5% PS (D, F) or 50% PS (E, G). I represents input, S represents supernatant, and P represents pellet. Final PLCβ concentration used in the experiments is 150 nM.

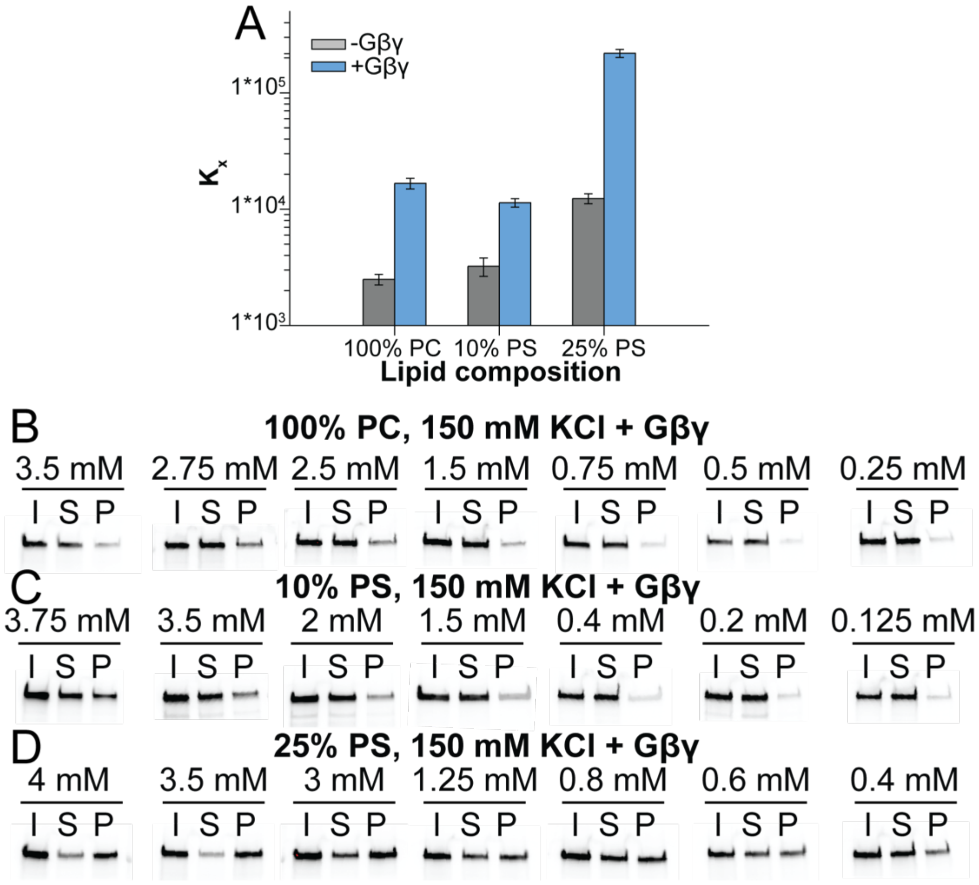


**Figure S5**: *Gβγ membrane recruitment efficiency depends on anionic lipid content*. **A**: Comparison of $K_x$ as a function of anionic lipid content in the absence (gray) or presence (blue) of Gβγ. Error bars are the error from the fit to determine $K_x$. **B-D**: Representative SDS-PAGE gels imaged for Cy5 fluorescence from partitioning experiments in the presence of Gβγ with 100% PC lipids (B), 10% PS (C), and 25% PS (D). I represents input, S represents supernatant, and P represents pellet. Final PLCβ concentration used in the experiments is 150 nM.
